# plantMASST - Community-driven chemotaxonomic digitization of plants

**DOI:** 10.1101/2024.05.13.593988

**Authors:** Paulo Wender P. Gomes, Helena Mannochio-Russo, Robin Schmid, Simone Zuffa, Tito Damiani, Luis-Manuel Quiros-Guerrero, Andrés Mauricio Caraballo-Rodríguez, Haoqi Nina Zhao, Heejung Yang, Shipei Xing, Vincent Charron-Lamoureux, Desnor N. Chigumba, Brian E. Sedio, Jonathan A. Myers, Pierre-Marie Allard, Thomas V. Harwood, Giselle Tamayo-Castillo, Kyo Bin Kang, Emmanuel Defossez, Hector H. F. Koolen, Milton Nascimento da Silva, Consuelo Yumiko Yoshioka e Silva, Sergio Rasmann, Tom W. N. Walker, Gaëtan Glauser, José Miguel Chaves-Fallas, Bruno David, Hyunwoo Kim, Kyu Hyeong Lee, Myeong Ji Kim, Won Jun Choi, Young-Sam Keum, Emilly J. S. P. de Lima, Lívia Soman de Medeiros, Giovana A. Bataglion, Emmanoel V. Costa, Felipe M. A. da Silva, Alice Rhelly V. Carvalho, José Diogo E. Reis, Sônia Pamplona, Eunah Jeong, Kyungha Lee, Geum Jin Kim, Yun-Seo Kil, Joo-Won Nam, Hyukjae Choi, Yoo Kyong Han, Si Young Park, Ki Yong Lee, Changling Hu, Yilun Dong, Shengmin Sang, Colin R. Morrison, Ricardo Moreira Borges, Andrew Magno Teixeira, Seo Yoon Lee, Bum Soo Lee, Se Yun Jeong, Ki Hyun Kim, Adriano Rutz, Arnaud Gaudry, Edouard Bruelhart, Iris F. Kappers, Rumyana Karlova, Mara Meisenburg, Roland Berdaguer, J. Sebastián Tello, David Henderson, Leslie Cayola, S. Joseph Wright, David N. Allen, Kristina J. Anderson-Teixeira, Jennifer L. Baltzer, James A. Lutz, Sean M. McMahon, Geoffrey G. Parker, John D. Parker, Trent R. Northen, Benjamin P. Bowen, Tomáš Pluskal, Justin J. J. van der Hooft, Jeremy J. Carver, Nuno Bandeira, Benjamin S. Pullman, Jean-Luc Wolfender, Roland D. Kersten, Mingxun Wang, Pieter C. Dorrestein

## Abstract

Understanding the distribution of hundreds of thousands of plant metabolites across the plant kingdom presents a challenge. To address this, we curated publicly available LC-MS/MS data from 19,075 plant extracts and developed the plantMASST reference database encompassing 246 botanical families, 1,469 genera, and 2,793 species. This taxonomically focused database facilitates the exploration of plant-derived molecules using tandem mass spectrometry (MS/MS) spectra. This tool will aid in drug discovery, biosynthesis, (chemo)taxonomy, and the evolutionary ecology of herbivore interactions.

## Main

Earth harbors around 450 plant families, 16,000 genera, and 350,000 vascular plant species^1^. Plants are essential to our planet’s health through their role in converting CO_2_ into oxygen and energy, sustaining animal life, including our own, and have been used by humans to treat diseases. Despite their ecological, nutritional, and medicinal value, the taxonomic distribution of plant metabolites is hard to establish. This is due to the absence of a common open-access spectral MS plant database and search engines. Furthermore, not every species distributed in the plant kingdom has been studied and not every imaginable metabolite has been discovered. Most chemotaxonomic studies of plants are limited to specific clades of the taxonomic tree^2–5^ and rely on data availability of known compounds in structural and spectral databases^6–14^, limiting the search of yet-to-be-characterized molecules. However, creating a chemophenetic plant metabolite inventory of all plant species found worldwide is within reach as mass spectrometry (MS) technologies and algorithms are continuously improving. To facilitate this, we have created plantMASST, a taxonomically-informed mass spectrometry search tool for plant metabolites within the Global Natural Products Social Molecular Networking (GNPS) ecosystem. plantMASST builds on the the approach taken for microbeMASST^15^, creating a digital inventory of untargeted plant metabolomics data. This enables the querying of MS/MS spectra corresponding to known and unknown molecules within a curated database of LC-MS/MS data from plant extracts, with results mapped across the plant taxonomic tree. As of May 2024, plantMASST curated reference datasets containing LC-MS/MS data from 19,075 plant extracts with over 100 million MS/MS spectra linked to their respective taxonomical information (**Figure 1a**). The plantMASST reference database results from community contributions and metadata curation from 90 scientists worldwide and it now includes 246 botanical families, 1,469 genera, and 2,793 species. To increase plantMASST coverage, we encourage the community to deposit new plant datasets in MassIVE (https://massive.ucsd.edu/) with associated metadata in the ReDU template^16^.

**Figure 1.**
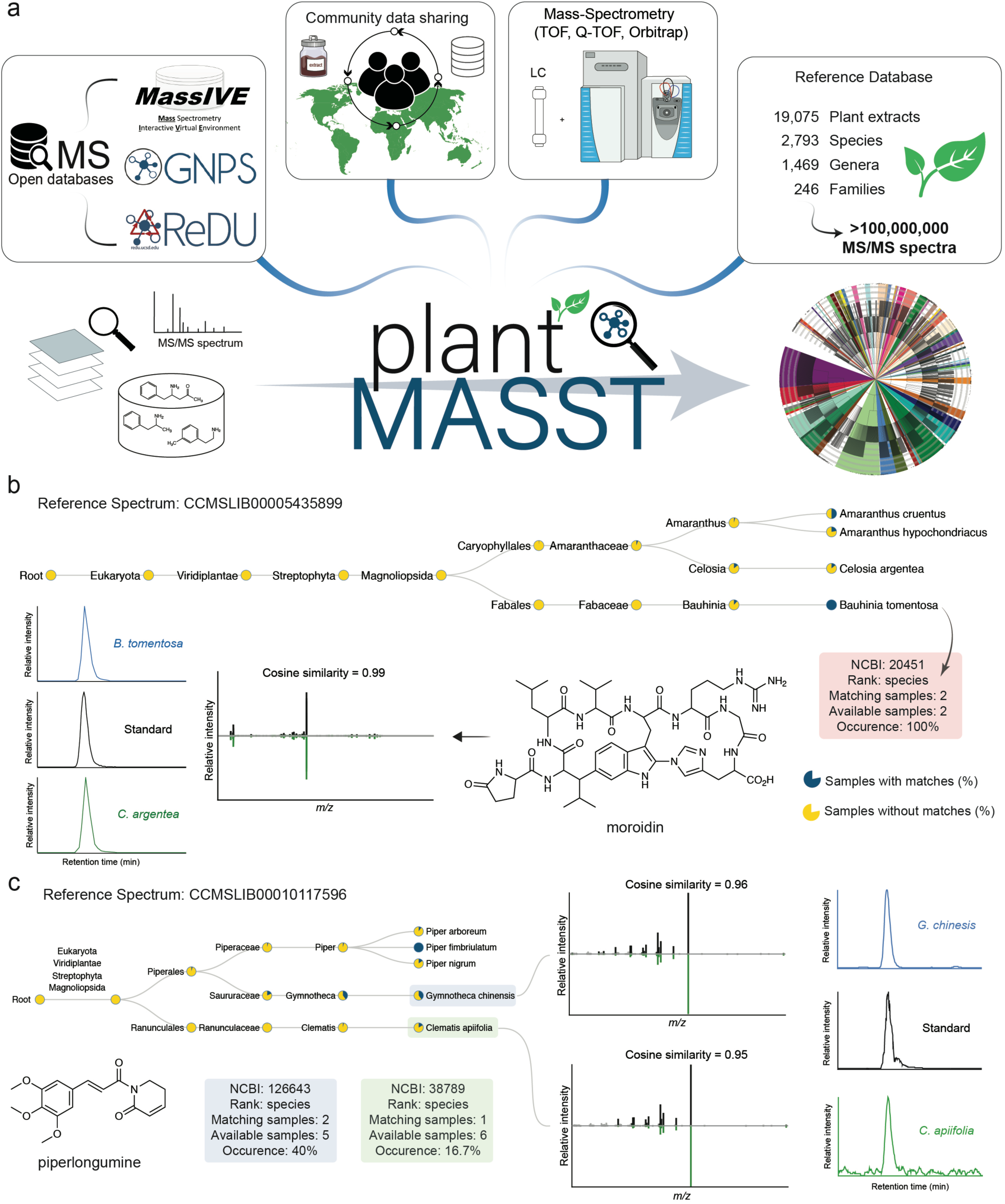
Schematic overview of plantMASST infrastructure and output. a) The creation of the plantMASST reference database involved utilizing 19,075 community-curated LC-MS/MS data and knowledge from MassIVE, GNPS^12^, and ReDU^16^. b) Example of plantMASST output interpretation of a specialized metabolite known to be produced by specific plants, moroidin (the molecule was confirmed using standard, **Supplementary Figure S4**). Pie charts display the percentage of matches within that taxonomic level detected against the plantMASST database. Blue indicates the percentage of samples with matches and yellow without matches. The reference MS/MS spectrum for moroidin (CCMSLIB00005435899) is available in the GNPS library. c) Output of piperlongumine (CCMSLIB00010117596) search. Mainly produced by *Piper* species, the molecule was also confirmed, for the first time, to be present in *Gymnotheca chinensis* and *Clematis apiifolia* leaf extracts via MS/MS and retention time matching to its commercial standard. In the mirror plots, the spectrum on the top is relative to piperlongumine, and the one on the bottom is a match to plantMASST. More tissues were also matched and the chromatograms can be found in **Supplementary Figure S5**.

A single MS/MS spectrum can be searched in plantMASST through its web interface (https://masst.gnps2.org/plantmasst/). The search output is a list of all data files where the queried MS/MS spectrum (within the user-defined scoring criteria) has been observed. Additionally, an interactive taxonomic tree of the results is generated, which can be easily explored (**Figure 1b,c**). The search is accomplished by providing either a Universal Spectrum Identifier (USI)^17^ or a precursor *m/z* and a corresponding list of fragment ions as *m/z*-intensity-pairs (**Supplementary Figure S1a**). Although the parameters are adjustable, the web interface provides the following default values: precursor and fragments mass tolerance set at 0.05 Da and cosine similarity of 0.7 with at least three matched fragment ions shared between the queried spectrum and the spectra available in plantMASST. “Analog search” can also be enabled by the user to explore MS/MS spectra from putative-related structural analogs or different ion forms of the same molecules (*e.g.*, different adducts, multimers, and in-source fragment ions). Moreover, plantMASST automatically performs spectral library searches against the reference libraries publicly available within the GNPS environment for direct metabolite annotation through MS/MS spectral matching. Building on the concepts of MASST^18^ and fastMASST^19^, plantMASST returns results within seconds and includes the taxonomic distribution of metabolites at once. The interactive taxonomic tree (and links to related tools in the GNPS ecosystem) used to visualize the results was built using NCBI^20^ taxonomy (**Supplementary Figure S1b**). Finally, users can explore matches inspecting the MS/MS mirror plots, with highlighted shared peaks, using the Metabolomics Spectrum Resolver^21^ and access the corresponding LC-MS/MS files using the GNPS Dashboard^22^.

Here, we highlight the use of plantMASST through three representative applications on natural product discovery and one about how to explore public human diet intervention datasets. First, researchers may desire to explore the taxonomic distribution of known plant-derived molecules. Therefore, as proof of concept, we investigated moroidin. This bicyclic octapeptide belongs to the BURP-domain-derived ribosomally synthesized and post-translationally modified peptides (RIPPs) family^23^, which was first isolated from *Dendrocnide moroides* Wedd. (Urticaceae)^24^ and *Celosia argentea* L. (Amaranthaceae)^25^. This molecule was described to induce apoptosis in the A549 human non-small cell lung cancer cell line and it has been gaining attention in the search for new drugs for cancer treatment^26^. We used plantMASST to search an MS/MS spectrum of the [M + 2H]^2+^ ion of moroidin and found four potential producer species across two different plant families, including yellow bauhinia (*Bauhinia tomentosa* Vell., Fabaceae) which was not a known producer of moroidin. Moroidin identification in the extract of *B. tomentosa* seeds was confirmed as Metabolite Identification level 1 by comparison to an authentic standard (**Figure 1b**) according to the Metabolomics Standard Initiative (MSI, **Supplementary Figure S4**)^27^. Another example of natural product discovery across the taxonomic domain is showcased by searching for piperlongumine, an amide alkaloid originally isolated from the *Piper* genus (Piperaceae), which has a reported antitumor activity^28^ (**Figure 1c**). This molecule is also currently being investigated for the treatment of glioblastoma^29^. Utilizing plantMASST, we observed that the piperlongumine MS/MS spectrum was also detected in two other botanical species: *Gymnotheca chinensis* Decne. (Saururaceae) and *Clematis apiifolia* DC. (Ranunculaceae), which has not been previously reported. MS/MS and retention time matching with a commercial standard allowed us to confirm the presence of piperlongumine in both species and achieve level 1 identification (**Figure 1c**)^27^. This expands the known natural reservoirs of piperlongumine beyond its primary source, *Piper longum* L. (Piperaceae), and highlights the versatility and efficacy of plantMASST in discovering pharmaceutically relevant phytochemicals in diverse plant families. It also suggests new research questions, such as why the production of these alkaloids is observed in such diverse plant species. Other plantMASST representative applications, as such caffeine, reserpine, icaridin, lutein, methoxsalen, and tryptophan, can be found in **Supplementary Figures S1, S3, S4,** and **S6.**

Beyond demonstrating the potential of plantMASST for highlighting specific plant metabolites with a narrow distribution within the plant kingdom, we built on recent observations that human neuroactives are also present in plants^30,31^. Many of them are involved in the signaling communication and adaptation of plants^31^. However, it is yet unknown to what extent they are distributed across the plant kingdom. As no report exists at the moment, we used plantMASST to search the MS/MS spectra of acetylcholine, dopamine, serotonin, glutamate, gamma-aminobutyric acid (GABA), and norepinephrine, all known neuroactives also produced by humans. Additionally, we searched for cannabidiol (CBD) and tetrahydrocannabinol (THC), two plant-derived metabolites known to affect human brain physiology for comparison (**Figure 2a,b**). Serotonin, also known as 5-HT, was detected in 61 out of 246 plant families available in plantMASST, with a notable prevalence in species belonging to the Malpighiaceae family, including *Banisteriopsis* genus (**Figure 2b**). This observation suggests this genus is a potential source of neuroactives such as serotonin. Additionally, it may also explain the neuroactive properties of species of this genus, such as *Banisteriopsis caapi* C.V. Morton (Malpighiaceae), which is used as the main ingredient of Ayahuasca, an indigenous beverage traditionally used in the Northwestern Amazon to treat mental health disorders^32^. These plants could be further evaluated for serotonin-mediated effects on breathing, sleep, arousal, and seizure control^33,34^. Other neuroactives were found in specific plant families. For instance, tryptamine was detected only in *Solanum lycopersicum* L. (Solanaceae); CBD and THC, two molecules that have neurological effects on humans, were found only in *Cannabis sativa* (Cannabaceae). Therefore, plantMASST can also enable users to explore the diversity of neuroactives in plants and highlight plants that could be further investigated for potential pharmacological applications or the extraction of bioactive compounds with medicinal properties.

**Figure 2.**
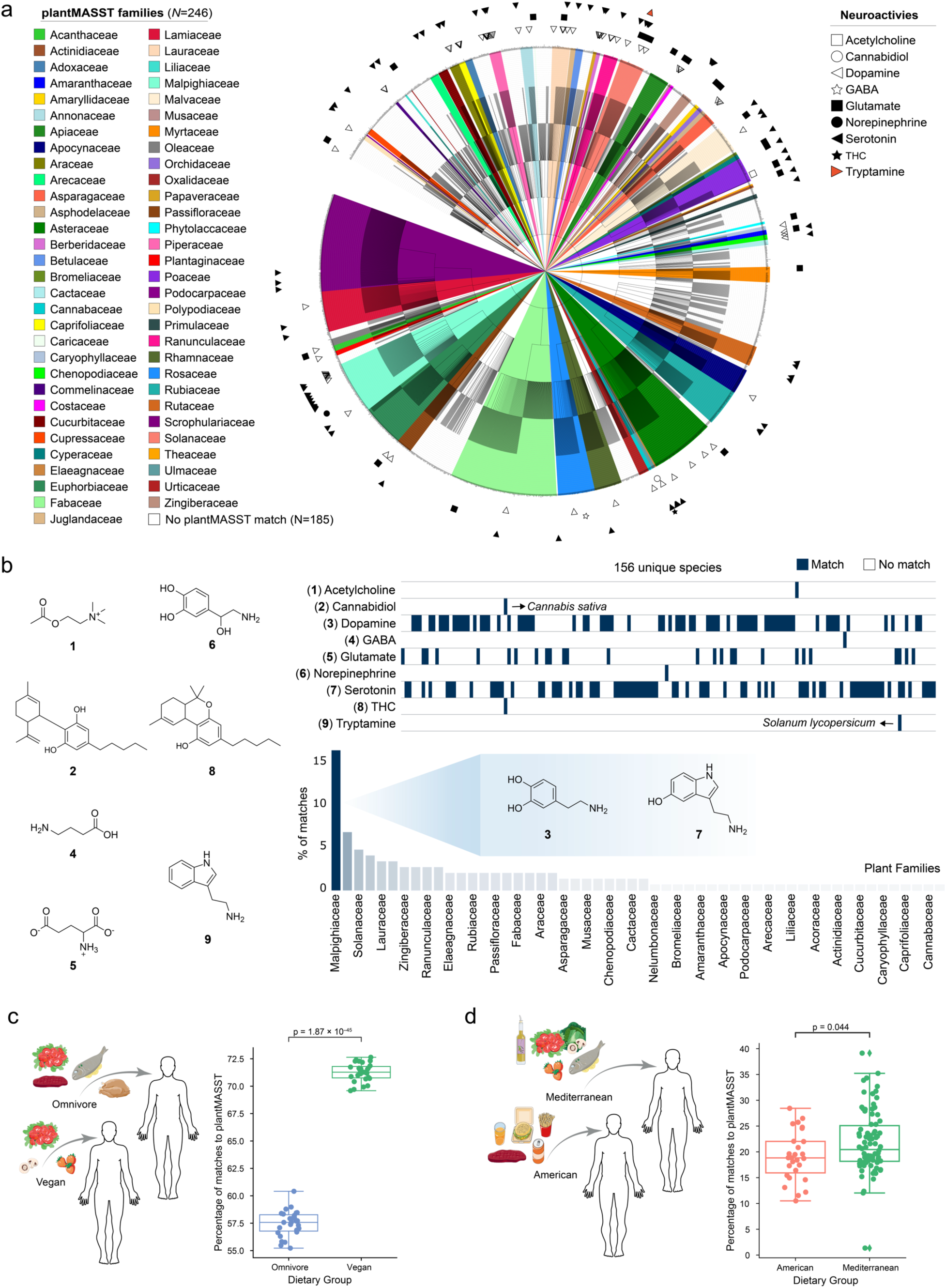
Use of plantMASST in the search for neuroactives in plants and plant-derived small molecules in public data sets of fecal samples of humans. a) Taxonomic distribution of neuroactive compounds across plant families. b) Heatmap showing the distribution of the nine neuroactives in 156 plant species. The bar plot highlights the sum of the percentage (%) of matches to serotonin/dopamine among the botanical families. c) Percentages of metabolites matched to plantMASST in a human publicly available diet-related dataset (GNPS/MassIVE: MSV000086989)^35,36^ containing fecal samples from vegans (n = 27) and omnivores (n = 27). d) Percentages of metabolites matched to plantMASST in a human diet-related dataset (GNPS/MassIVE: MSV000093005)^37^ containing fecal samples from patients consuming either an American (n = 27) or Mediterranean (n = 82) diet. Boxplots represent first (lower), median, and third (upper) quartiles. Upper and lower whiskers extend to the closest value to +/- 1.5 * interquartile range (IQR). The independent two-sided t-test was applied in Fig 2c, while the two-sided Mann-Whitney-Wilcoxon test was applied in Fig 2d, and statistical significance was observed for p < 0.05. Food icons were obtained from Bioicons.com.

Although plantMASST and the underlying reference database can be leveraged in many ways, we showcase a last example of how plantMASST can be used to detect human dietary consumption of plants. We reanalyzed all the MS/MS spectra from two public metabolomics datasets of human fecal data of diet-related studies; one comparing vegan *vs* omnivore diets^35,36^, and the other American *vs* Mediterranean diets^37^. We observed a higher percentage of MS/MS matches to plant metabolites in the vegan group when compared to the omnivore group (**Figure 2c**). Interestingly, also the subjects consuming the Mediterranean diet had more MS/MS matches to plant metabolites compared to the American diet (**Figure 2d**). These results suggest that plantMASST could also be used to define the nature of plant-derived dietary patterns.

It is important to bear in mind certain limitations when interpreting the results of plantMASST since the current reference database contains diverse experimental and acquisition conditions. This includes the plant organ, plant growth conditions, the biome of origin, extraction methods, sample preparation, instrument geometries, and defined collision energies, among others. Therefore, even if the plant is known to be a producer of a compound of interest, a plantMASST match might be missed because of fewer fragment ion matches to low-intensity MS/MS ions or because no MS/MS of the compound was triggered due to low precursor ion intensity. If the users decrease the tolerances of minimum MS/MS fragment peaks to find more matches, it may result in more false positive matches. Further, although many MS/MS spectra will be quite specific to one molecule (*e.g.*, moroidin and reserpine, **Supplementary Figure S2a**), isomers that have the same mass, can have nearly identical MS/MS spectrum, which is the case of quercetin and morin (**Supplementary Figure S2b**). This means that in some cases the taxonomic interpretation of a family of molecules may not map well to the individual molecules. Finally, there are currently only 2,793 species of plants, representing a fraction of the species that exist. Despite these challenges, plantMASST represents an important advance in our ability to map the plant taxonomic distribution of metabolites, especially as researchers continue to expand the taxonomic curation of untargeted plant metabolomics data. We expect that plantMASST will potentially have profound implications for the fields of drug discovery, nutrition, and chemical ecology.

## Methods

### Data collection and curation

To enable the taxonomic search of known and unknown compounds and to make a dent in capturing chemotaxonomic data from all available plant extracts, 211 publicly available MS/MS datasets in the GNPS/MassIVE were manually compiled and each file, within these datasets, was taxonomically defined possibly to species level. For the samples in which the species was not known, the closest known taxonomic rank was defined (e.g., genus or family). These datasets consisted of 20,209 unique LC-MS/MS files, representing plant extracts inclusive of all plant tissue types (leaves, stems, seeds, among others) and habits (trees, shrubs, lianas, herbs, etc). To collect this number of samples, we made an open call to the scientific community to deposit plant-related datasets in GNPS/MassIVE, which led to the deposition of an additional 25 datasets between December 2022 and March 2023, resulting from the efforts of 12 research groups across the world. All the collected information of each file part of plantMASST is available on GitHub and contains the following information: the path of each file, the filename in the format ‘Dataset/Filename’, the MassIVE ID, the taxon name, the NCBI Taxonomy ID, ReDU availability, whether the file is relative to a blank or a QC, and the USI of the file.

To get the NCBI taxonomy IDs, the datasets containing ReDU metadata were matched to NCBI IDs. When only a table containing the sample names was provided, the NCBI taxonomy IDs were manually retrieved from the NCBI Taxonomy web browser (https://www.ncbi.nlm.nih.gov/taxonomy).

### plantMASST taxonomic tree generation

The plantMASST taxonomy tree was created with R Studio 4.2.2 and Python 3.10. Only distinct NCBI IDs (n = 3,173) were kept in the database. To obtain the complete lineage of every NCBI ID^38^, the ‘taxize’ package (v.0.9.100) was used, namely its categorization function. The main taxonomic ranks (kingdom to species) along with subgenus, subspecies, and variations were maintained to create taxonomic trees with an equal number of taxa nodes. After importing the NCBI ID list for every taxon into Python, a taxonomic tree was created using the ETE toolkit^39^. After that, the created Newick tree was transformed into JSON format, and details like taxonomic rank and number of accessible samples were added. All taxonomic entries in plantMASST are classified according to the NCBI taxonomy^20^, which includes 246 families, 1,469 genera, and 2,793 species. Thus, while plantMASST encompasses more than 50% of the world’s botanical families present in the current community curation, less than 1% of species are represented. The representation of the plant data will continue to grow as the scientific community continues to contribute to expanding publicly available plant-related datasets.

### MASST searching

plantMASST can be used as a web application by accessing https://masst.gnps2.org/plantmasst/. This web application was built using Dash and Flask open-source libraries for Python (https://github.com/mwang87/GNPS_MASST/blob/master/dash_plantmasst.py). Searches in the web interface can be done by inputting a Spectrum USI or the spectrum peaks of an MS/MS spectrum and further clicking on ‘Search plantMASST by USI’ or ‘Search plantMASST by Spectrum Peaks’. The default search parameters are cosine threshold: 0.7; minimum matched peaks: 3; precursor mass and fragment tolerance: 0.05 Da; analog search: no. Users familiar with Python can also create batch searches via the command line, which produces results for plantMASST, microbeMASST, and foodMASST. Details can be found at https://zenodo.org/records/10909241. With this approach, users have to provide either a CSV or TSV file containing the USIs to be searched or an MGF file containing precursors and MS/MS spectra of the ions. The same settings described in the API web interface such as minimum cosine, *m/z* tolerance, precursor *m/z*, and minimum fragments matched, are adjustable. To create the resulting taxonomic tree, the JSON file of the complete plantMASST taxonomic tree is filtered and converted into a D3 JavaScript object that can be visualized as an HTML file.

### plantMASST applications

plantMASST can be used in a variety of scenarios. First, we showed the taxonomic distribution of specific metabolites present in the GNPS public libraries (moroidin, caffeine, reserpine, icaridin, lutein, methoxsalen, and piperlongumine). These searches were done using the web interface and the default parameters. Second, selected neuroactives with MS/MS spectra available in the GNPS public libraries (acetylcholine, dopamine, GABA, glutamate, norepinephrine, serotonin, CBD, THC, and tryptamine) were searched against plantMASST with the web interface using the default parameters. The matches obtained were manually inspected via mirror plots and the low-quality matches were filtered out. The tables containing the taxonomic distribution were downloaded and combined to visualize them using the Interactive Tree of Life (iTOL)^40^. A heatmap was generated in Python (version 3.11) to show the plantMASST matches of these neuroactives to different plant species. The packages ‘pandas’ (version 2.2.2) and ‘plotly’ (version 5.21.0) were used for this analysis. Finally, we re-analyzed two diet-related MS/MS studies (vegan vs omnivore diet: MSV000086989; American vs Mediterranean diet: MSV000093005). The raw data was processed in MZmine3 (version 3.4.27). The MZmine3 batch files used in each study are available on GitHub (https://github.com/helenamrusso/plantmasst), in addition to the generated output files. The generated .mgf files were used as input for the plantMASST batch search (https://github.com/robinschmid/microbe_masst) using the following parameters: cosine threshold: 0.7; minimum matched peaks: 4; precursor mass and fragment tolerance: 0.02 Da; analog search: off. Boxplots showing the percentages of matches to plantMASST in each sample were generated in Python using the ‘seaborn’ package (version 0.11.2). The Shapiro-Wilk test was used to assess the normality of the data.

### LC-MS/MS analysis

#### Retention time matching for moroidin

Moroidin was isolated and purified as described^23^ from *Celosia argentea* flower and seed material. All materials were purchased from Fisher Scientific unless otherwise noted. *Celosia argentea* seeds were purchased from Seedville^TM^, and *Bauhinia tomentosa* seeds were purchased from rarepalmseeds.com^TM^. 0.2 g of *C. argentea* seeds or *B. tomentosa* seeds were each ground with mortar and pestle, extracted with 3 mL methanol for 1 h shaking at 200 rpm at 37 °C in a 7 mL glass scintillation vial (Fisher Scientific 03-337-26). The methanol extracts were dried under nitrogen gas and resuspended in 3 mL of deionized water. The resuspensions were partitioned twice with hexane (1:1, v/v), partitioned twice with ethyl acetate (1:1, v/v), and extracted once with 3 mL n-butanol. The n-butanol extract was dried *in vacuo* in a Thermo Scientific SPD140P1 speed vac and resuspended in 2 ml of 80% methanol for LC-MS/MS analysis. The moroidin standard had a final concentration of 500 nM in 80% methanol.

LC-MS/MS analysis for moroidin analysis was carried out on a Thermo H-ESI-Q-Exactive Orbitrap mass spectrometer coupled to a Thermo Vanquish ultra-HPLC system. H-ESI parameters were as follows: spray voltage: +4.2 kV; ion transfer tube temperature: 320 °C; S-lens RF: 50 (arb units); sheath gas flow rate: 35 (arb units); Sweep Gas: 0, and auxiliary gas flow rate: 3 (arb. units). The LC-MS/MS settings were as follows: injection volume 5 µl; LC, Phenomenex Kinetex 2.6 μm C18 reverse phase 100 Å 150 ×3 mm LC column; LC gradient, solvent A, 0.1% formic acid; solvent B, acetonitrile (0.1% formic acid); 0 min, 10% B; 5 min, 60% B; 5.1 min, 95% B; 6 min, 95% B; 6.1 min, 10% B; 9.9 min, 10% B; 0.5 ml min^−1^; MS, positive ion mode; full MS, resolution 70,000; mass range 400–1,200 *m/z*; dd-MS^2^ (data-dependent MS/MS), resolution 17,500; AGC target 1 × 10^5^, loop count 5, isolation width 1 *m/z*, collision energy 25 eV and dynamic exclusion 0.5 s. LC-MS/MS data were analyzed with QualBrowser in the Thermo Xcalibur software package (v4.3.73.11).

#### Retention time matching for piperlongumine

Piperlongumine was obtained as a commercial standard (>97 % HPLC quality, SML0021, Sigma-Aldrich) to enable the confirmation of the presence of this compound in additional plant extracts. LCMS grade acetonitrile, water, and formic acid (all Optima^®^ LCMS Grade, Fisher Scientific, Germany) were used for the analyses. The Pierre Fabre Laboratories supplied the dried extracts for the *Gymnotheca chinensis* leaves (V113160) and green stems (V113161), *Clematis apiifolia* leaves (V113131), roots (V113132) and green stems (V113133). The extracts were prepared as described by Allard *et al.* (2023)^41^. All the samples were dissolved in methanol at a concentration of 5 mg/mL. The piperlongumine standard was prepared at a concentration of 100 µg/mL in methanol.

Analyses were performed with a Vanquish Horizon (Thermo Scientific) equipped with a binary pump H, a dual split sampler HT and a Diode Array detector FG coupled to an Orbitrap Exploris 120 mass spectrometer (Thermo Scientific), and a Corona Veo RS Charged Aerosol Detector (CAD, Thermo Scientific). The Orbitrap employs a heated electrospray ionization source (H-ESI) with the following parameters: spray voltage: +3.5 kV; ion transfer tube temperature: 320.00 °C; vaporizer temperature: 320.00 °C; S-lens RF: 45 (arb units); sheath gas flow rate: 35.00 (arb units); Sweep Gas (arb): 1, and auxiliary gas flow rate: 10.00 (arb. units).

The mass analyzer was calibrated using a mixture of caffeine, methionine-arginine-phenylalanine-alanine-acetate (MRFA), sodium dodecyl sulfate, sodium taurocholate, and Ultramark 1621 in an acetonitrile/methanol/water solution containing 1% formic acid by direct injection. Control of the instruments was done using Thermo Scientific Xcalibur software v. 4.6.67.17. Full scans were acquired at a resolution of 30,000 FWHM (at *m/z* 200) and MS^2^ scans at 15000 FWHM in the range of 100−1000 *m/z*, with 1 microscan, time (ms): 200 ms, an RF lens (%): 70; AGC target custom (Normalized AGC target (%): 300); maximum injection time (ms): 130; Microscans: 1; data type: profile; Usue EASY-IC(TM): ON. The Dynamic exclusion mode: Custom; Exclude after n times: 1; Exclusion duration (s): 5; Mass tolerance: ppm; low: 10, high: 10, Exclude isotopes: true. Appex detention: Desired Apex Window (%): 50. Isotope Exclusion: Assigned and unassigned with an exclusion window (*m/z*) for unassigned isotopes: 8. The Intensity threshold was set to 2.5E^5^ and a targeted mass exclusion list was used. The centroid data-dependent MS^2^ (dd-MS^2^) scan acquisition events were performed in discovery mode, triggered by Apex detection with a trigger detection (%) of 300 with a maximum injection time of 120 ms, performing 1 microscan. The top 3 abundant precursors (charge states 1 and 2) within an isolation window of 1.2 *m/z* were considered for MS/MS analysis. For precursor fragmentation in the HCD mode, a normalized collision energy of 15, 30, and 45% was used. Data was recorded in profile mode (Use EASY-IC(TM): ON).

The chromatographic separation was done on a Waters BEH C18 column (100 × 2.1 mm i.d., 1.7 μm, Waters, Milford, MA) using a gradient as follows (time (min), %B):0.00, 2; 3.1, 2; 17.36, 99; 23.12, 99; 23.60, 2; 26.00, 2. The mobile phases were (A) water with 0.1% formic acid and (B) acetonitrile with 0.1% formic acid. The flow rate was set to 500 μL/min, the injection volume was 3 μL, and the column was kept at 40 °C. The PDA detector was used from 210 to 400 nm with a resolution of 1.2 nm. The CAD was kept at 40 °C, 5 bar N_2_, and power function 1 for a data collection rate of 20 Hz.

## Acknowledgments

This research was partly supported by NIH R01GM107550 (PCD) and R01DK131753 (SS) and USDA grant 2018-67001-28265 (SS). TRN, BPB, and TVH are supported by the U.S. Department of Energy Joint Genome Institute (https://ror.org/04xm1d337), a DOE Office of Science User Facility, supported by the Office of Science of the U.S. Department of Energy operated under Contract No. DE-AC02-05CH11231. MW was supported by UCR Startup, NIH 5U24DK133658-02, and the U.S. Department of Energy Joint Genome Institute (https://ror.org/04xm1d337), a DOE Office of Science User Facility, is supported by the Office of Science of the U.S. Department of Energy operated under Contract No. DE-AC02-05CH11231. Data from the Madidi Project in Bolivia were supported by grants from: the Living Earth Collaborative at Washington University in St. Louis to JAM, BES, DH, and JST; the US National Science Foundation (DEB 0101775, DEB 0743457, DEB 1836353), and the National Geographic Society (NGS7754-04 and NGS 8047-06). Data from the Forest Global Earth Observatory (ForestGEO) were supported by a grant from Corteva Agrisciences to BES and SJW and by US National Science Foundation (NSF) DEB grants 2240430 and 2240431to BES and JAM. KBK, EJ, and KL were supported by the National Research Foundation of Korea (NRF) grants funded by the Korean Government (MSIT) (NRF-2020R1C1C1004046, 2022R1A5A2021216, and 2022M3H9A2082952). GJK and HC were supported by the National Research Foundation of Korea (NRF) grants funded by the Korean Government (MSIT) (Grant No. NRF-2020R1A6A1A03044512, and NRF-2021R1A2C1010727). RB and RK are supported by the Dutch Research Council (NWO/OCW), as part of the MiCRop Consortium Programme, Harnessing the second genome of plants (grant number 024.004.014). MM and IFK were supported by the Dutch Research Council (NWO/ALWGR.2017.004). SR and ED were supported by the Swiss National Science Foundation (grant number 204811 and 179481). GG was supported by the Swiss National Science Foundation (grant number 183365). PMA and EB were supported by a Swiss Open Research Data Grant (CHORD) in Open Science I from swissuniversities. PMA and EB are thankful to the Botanical Garden of the University of Fribourg team for access to the plant material profiled in MSV000090090. J-L.W. and L-M.Q-G are thankful to the Pierre-Frabe Laboratories for access to their plant collection profiled in sets MSV00008772 and MSV000087970, and for the support of the Swiss National Science Foundation (SNF N ◦ CRSII5_189921/1). The Canada Research Chairs program and a Natural Science and Engineering Research Council Discovery Grant supported JLB. JAL was supported by the Utah Agricultural Experiment Station (1398 and 1711). HHFK, GAB, EVC, FMAS, and EJSPL were supported by the Amazonas State Research Support Foundation (FAPEAM) and National Council for Scientific and Technological Development (CNPq grants: 306335/2023-9 and 305602/2023-3). HHFK was supported by Financiadora de Estudos e Projetos (FINEP) grant no. 01.22.0401.00. LSM was supported by the São Paulo Research Foundation, (FAPESP, grant 20/08270-0). The Gordon and Betty Moore Foundation also supported A.M.C.-R. and P.C.D. grant GBMF12120. HY was supported by the National Research Foundation of Korea (NRF) grants funded by the Korean Government (MSIT) (NRF-2021R1C1C1011857). KYL was supported by the National Research Foundation of Korea (NRF) grants funded by the Korean Government (MSIT) (NRF-2019R1A6A1A03031807 and 2021R1A2C1093814). KHK, SYL, BSL, and SYJ were supported by the National Research Foundation of Korea (NRF) grants funded by the Korean government (MSIT) (2019R1A5A2027340 and 2021R1A2C2007937). JDP was supported by the US National Science Foundation DEB Award No. 2044406. NB was partially supported by NIH award R24GM148372. GT-C and JMC-F were partially supported by the Research Vicerectory of the University of Costa Rica, grant 809-B8A15.

## Author contributions

P.W.P.G., H.M-R., and P.C.D. designed the project.

S.Z., R.S., B.S.P., N.B. and M.W. developed the code for the plantMASST interface.

P.W.P.G., H.M-R., and T.D. tested the website and prepared examples from public data.

A.M.C-R., S.X., H.Y., D.N.C., B.E.S., J.A.M., P-M.A., J-L.W., G.T-C., K.B.K., T.D., E.D., H.H.F.K., M.N.S., C.Y.Y.S., S.R., T.W.N.W., G.G., L-M.Q-G., J.M.C-F., B.D., H.K., K.H.L., M.J.K., W.J.C., Y-S.K., E.J.S.P.L., L.S.M., G.A.B., E.V.C., F.M.A.S., A.R.V.C., J.D.E.R., S.P., E.J., K.L., G.J.K., Y-S.K., J-W.N., H.C., Y.K.H., S.Y.P., K.Y.L., C.H., Y.D., S.S., C.R.M., R.M.B., A.M.T., D.E.L., B.S.L., S.Y.J., K.H.K., A.R., A.G., E.B., I.F.K., R.K., M.M., R.B., S.T., D.H., A.F.F., L.C., J.W., D.N.A., K.J.A-T., J.L.B., C.W.D., J.A.L., S.M.M., G.G.P., J.D.P., T.P., J.J.J.v.d.H., L-F.N., N.B., J.C., R.D.K., contributed with untargeted LC-MS/MS data published at MassIVE.

P.W.P.G., H.M-R., and P.C.D. wrote the article, and all authors reviewed and edited the text and figures.

## Data availability

Data used to generate the reference database of plantMASST are publicly available at GNPS/MassIVE (https://massive.ucsd.edu/). A list with all the accession numbers (MassIVE IDs) of the studies used to generate this tool is available on GitHub (https://github.com/helenamrusso/plantmasst, plant_masst_table.csv). All the taxonomic trees shown in this manuscript can be interactively explored by downloading the .html files available on GitHub (https://github.com/helenamrusso/plantmasst). To help interpret and establish that distinct plant species’ small molecules were only found, known molecules already present in the GNPS library (https://library.gnps2.org/) were employed.

- Moroidin (CCMSLIB00005435737)
- Piperlongumine (CCMSLIB00010117596)
- Caffeine (CCMSLIB00006365672)
- Quercetin (CCMSLIB00010118464)
- Morin (CCMSLIB00010122829)
- Reserpine (CCMSLIB00010110971)
- Icaridin (CCMSLIB00000565057)
- Lutein (CCMSLIB00005777353)
- Methoxsalen (CCMSLIB00006417040)
- Cannabidiol (CCMSLIB00009943776)
- Tryptophan (CCMSLIB00003136269)
- Acetylcholine (CCMSLIB00000578035)
- Dopamine (CCMSLIB00006121682)
- GABA (CCMSLIB00000215050)
- Glutamate (CCMSLIB00000081783)
- Norepinephrine (CCMSLIB00000219763)
- Serotonin (CCMSLIB00006114036)
- THC (CCMSLIB00005774204)
- Tryptamine (CCMSLIB00004693658)

Data used to search for plant-derived molecules (**Figure 2c**) from fecal samples of vegans and omnivores is publicly available in GNPS/MassIVE under the accession number MSV000086989. Data used to assess plant-derived molecules in fecal samples from people subjected to an American and Mediterranean diet is publicly available in GNPS/MassIVE under the accession number MSV000093005. Data acquired for retention time matching between piperlongumine standard and plant extracts is available in GNPS/MassIVE under the accession number MSV000094562.

## Code availability

The plantMASST web application is accessible at https://masst.gnps2.org/plantmasst/, and the open-source code is available under MIT license on GitHub (https://github.com/robinschmid/microbe_masst). The code used to generate the plantMASST web interface can be accessed on GitHub (https://github.com/mwang87/GNPS_MASST). The code used to perform the analyses and generate the figures present in the manuscript can be downloaded from GitHub (https://github.com/helenamrusso/plantmasst).

## Supplementary Figure

Supplementary Figure S1. plantMASST web app; Supplementary Figure S2. High-resolution MS/MS spectra; Supplementary Figure S3. Complementary output of plantMASST; Supplementary Figure S4. plantMASST examples of four rarely occurring plant-derived drugs; Supplementary Figure S5. Retention time matching with piperlongumine; Supplementary Figure S6. Taxonomic tree of tryptophan.

## Supplementary information

### Complementary use cases

In the plantMASST web interface, a single search was performed using GNPS reference MS/MS spectra of caffeine, and the taxonomic tree was generated as a result (**Supplementary Figure S1b)**. Caffeine is a drug found in many plants; it is a central nervous system stimulant, enhancing wakefulness, improving alertness, and decreasing weariness^42^. The output searches from plantMASST show caffeine was most notably detected in species used to prepare these beverages. For instance, *Camellia sinensis* (Theaceae) is traditionally used for caffeinated teas^43^. Our findings also support caffeine content in cacao (*Theobroma cacao*, Malvaceae), and *Coffea arabica* (Rubiaceae, from which coffee beans are created). Moreover, searching with rare metabolites, such as reserpine was detected in 0.010% of the samples (**Supplementary Figure S3**). Reserpine is a drug that is used clinically to treat illnesses such as hypertension and Parkinson’s disease^44^. It was isolated for the first time from the plant *Rauvolfia* (=*Rauwolfia) serpentina* (Apocynaceae) and plays a role by blocking noradrenaline, dopamine, and serotonin reuptake in nerve cells, resulting in relaxing and blood pressure-lowering effects^45^. Reserpine is an example of a natural compound found in plants with medicinal effects, and in this correspondence, given the ability of plantMASST reserpine is being putatively described for the first time in *R. media*, a species from the same genus of *R. serpentina*.

Another utility of plantMASST is for searching for plants that can produce drugs with diverse chemical backbones. Specifically, we examined four approved plant-derived drugs, icaridin, lutein, methoxsalen, and cannabidiol, from the Drugbank database and observed their presence in taxonomically distinct species (**Supplementary Figure S4**). Icaridin is known due to its plant insect-repellent properties. Also, it is often used topically as an insect repellent to protect against mosquito bites, ticks, and other insects^46^. Otherwise, lutein is a naturally occurring carotenoid pigment found in various plants and vegetables, and it is known for its role in eye health^47,48^. It is a major component of the macular pigment in the human retina and is believed to help protect the eyes from oxidative damage and age-related macular degeneration^48^. Methoxsalen is a naturally occurring compound found in certain plants, including citrus fruits like oranges and lemons, as well as in various plants such as figs and parsley^49^. Methoxsalen is primarily used in a medical procedure known as PUVA (psoralen plus ultraviolet A) therapy, and it is also used to treat skin conditions like psoriasis, vitiligo, and eczema^50^.

Last, cannabidiol (CBD) is one of the many compounds derived from the resin of cannabis flowers and can be extracted for various uses^51^. CBD has gained significant attention for its potential medicinal benefits. It is used in the treatment of epilepsy (specifically, Dravet syndrome and Lennox-Gastaut syndrome), chronic pain management, anxiety, and as an anti-inflammatory agent^52–54^. Thus, plantMASST allowed us to find these drugs in different families *e.g.*, icaridin was detected in species from several families. Otherwise, the other drugs were mainly detected in specific species, for instance, lutein in *Eremophila calcicola* (Scrophulariaceae), methoxsalen in *Citrus* (Rutaceae), and CBD in *Cannabis sativa* (Cannabaceae). Therefore, these findings indicated that the plantMASST can be leveraged to hypothesize the discovery of abundant sources of plants containing structurally analogous drugs and their potential origins of these drugs/MS-MS signals from a database containing metabolome information from extracts of plants. In other cases, such as tryptophan, it is very frequently detected (18.69% of all the samples, **Supplementary Figure S6**), highlighting that plantMASST can be leveraged to hypothesize if the MS/MS is a taxonomically common or rare metabolite.

**Supplementary Figure S1.**
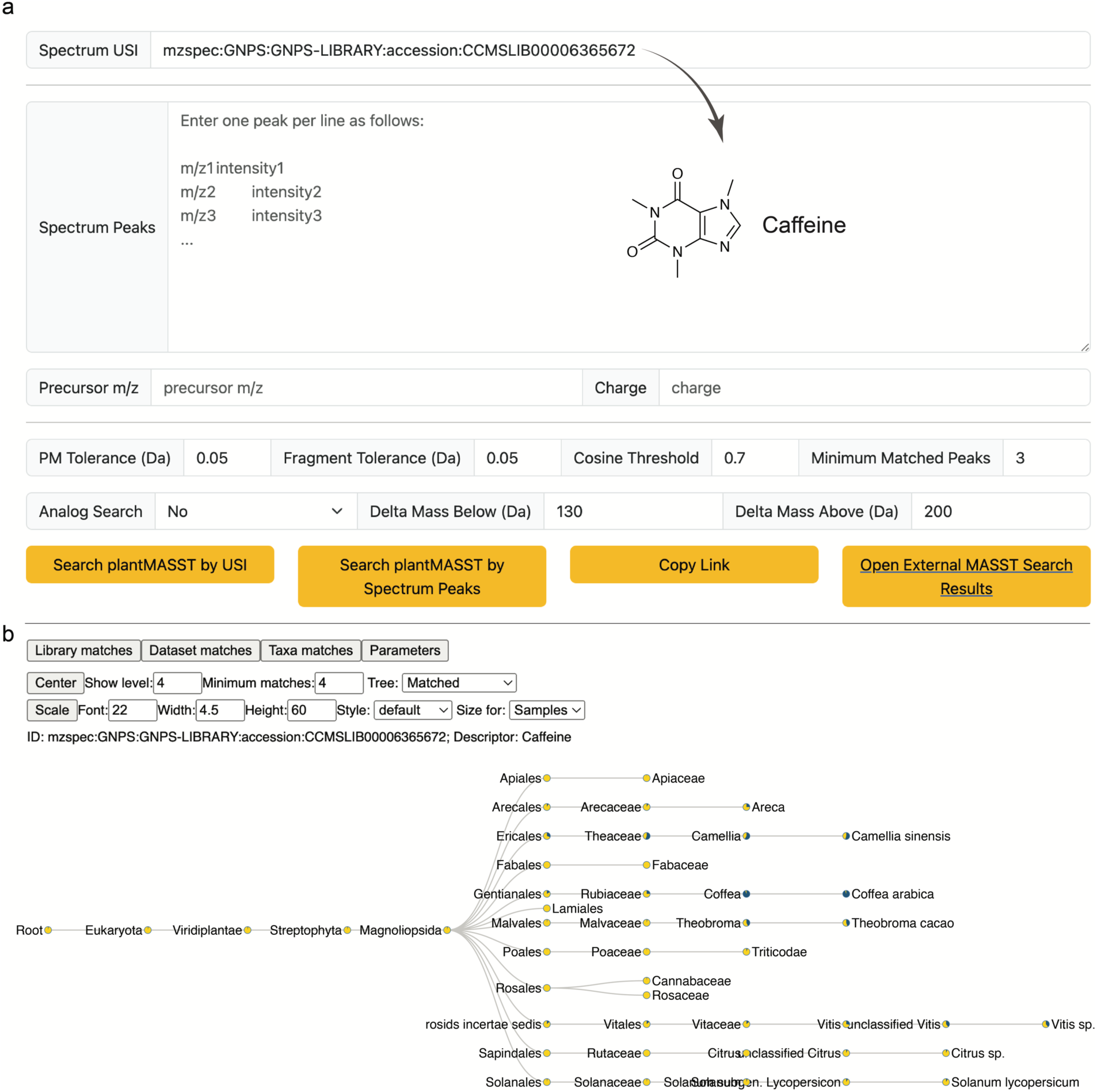
plantMASST web app. a) The plantMASST web app is available at https://masst.gnps2.org/plantmasst/. People can search the plantMASST reference database for MS/MS spectra by entering a USI in ‘Spectrum USI’ or fragment ions, intensities, and precursor mass in ‘Spectrum Peak’ and ‘Precursor *m/z*’, respectively. Precursor and fragment ion tolerances, cosine threshold, and minimum matching peaks are adjustable. Analog search is also available. Finally, based on the information presented, users can enter a search query by clicking either ‘Search plantMASST by USI’ or ‘Search plantMASST by Spectrum Peak’. Search jobs can be readily shared by clicking on the ‘Copy Link’ button. Reference MS/MS spectrum (CCMSLIB00006365672) available in the GNPS library was used to search for caffeine in the plantMASST database. b) interactive taxonomic tree and distribution of caffeine in plant species according to the NCBI taxonomy. Also, links to related tools in the GNPS ecosystem are provided (library, dataset, and taxa matches). These matches represent level 2 annotations according to the Metabolomics Standards Initiative^27^.

**Supplementary Figure S2.**
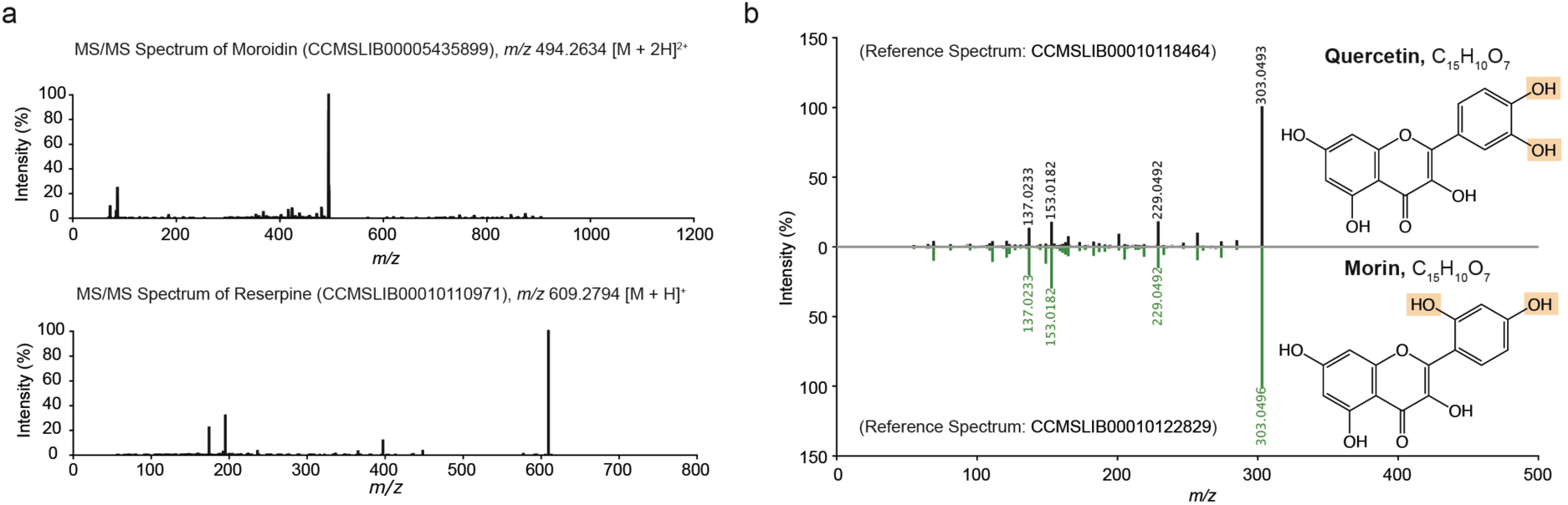
High-resolution MS/MS spectra of moroidin and reserpine. a) Single spectrum of moroidin and reserpine. b) Mirror plot between the MS/MS spectra of quercetin and morin. Although they have the same spectrum, their structures are different.

**Supplementary Figure S3.**
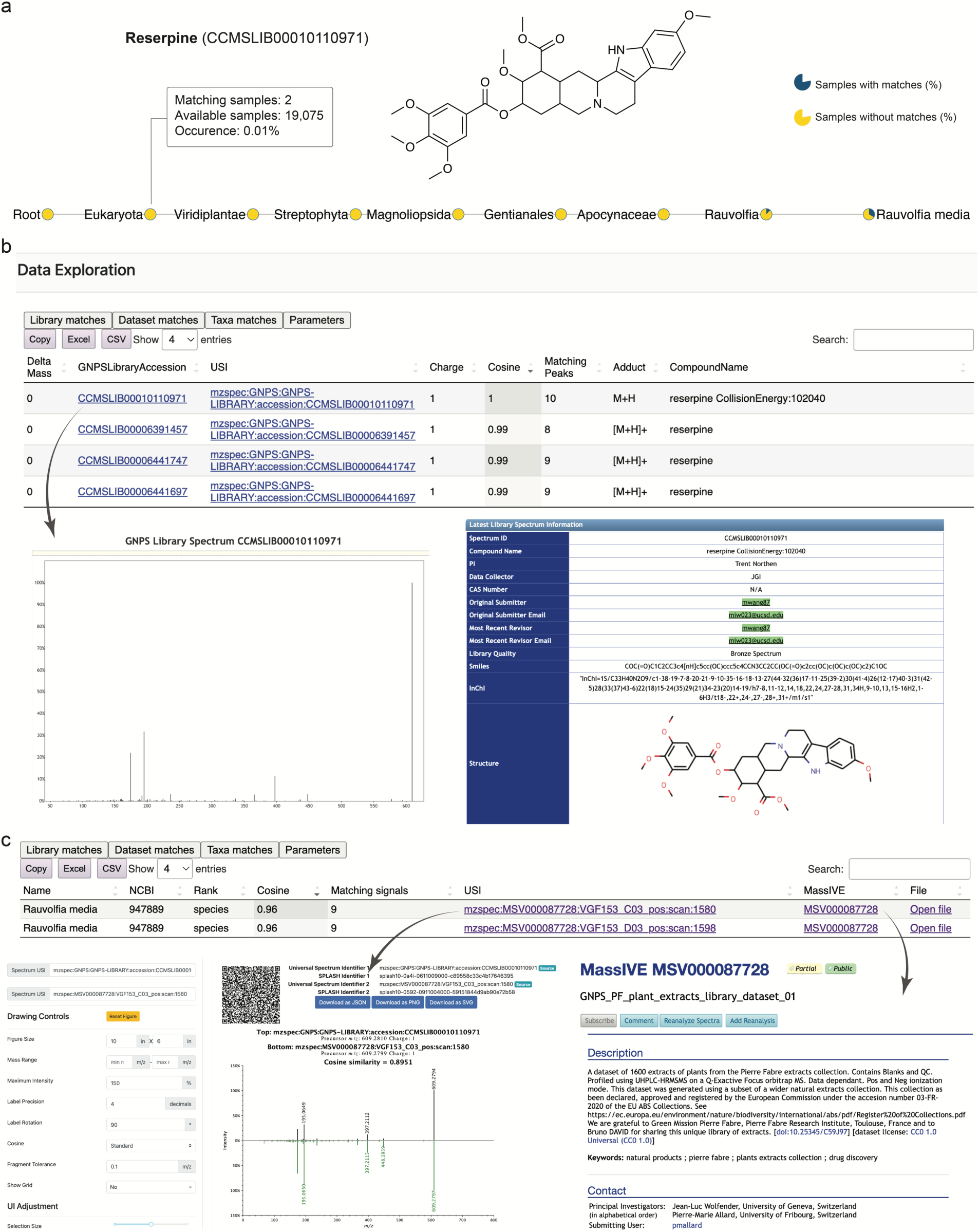
Complementary output of plantMASST. a) Example of plantMASST taxonomic tree of reserpine, known to be produced by specific plants. The taxonomic tree shows the MS/MS spectrum of reserpine searched against the plantMASST database was detected only in *Ochrosia elliptica.* It represents 0.010% of the samples available in the database (2 samples out of 19,075). Pie charts display the percentage of matches within that taxonomic level detected against the plantMASST database. Blue indicates the percentage of samples with matches and yellow without matches. Reference MS/MS spectra of reserpine (CCMSLIB00010110971), and it represents a level 2 annotation according to the Metabolomics Standards Initiative^27^. b) The MS/MS spectrum is searched against the GNPS libraries, and possible annotations are returned if matches are identified. Users can visit the accompanying GNPS Library Spectrum page for information on the reference spectrum. c) Data on matched scans in the sample from various taxa is supplied. Furthermore, users can visualize the mirror plot between the queried spectrum and spectrum from datasets included in the plantMASST database, such as the similarity score, and matching fragments. The user can also obtain the project’s MassIVE accession number as well as contact information for the person who contributed the data.

**Supplementary Figure S4.**
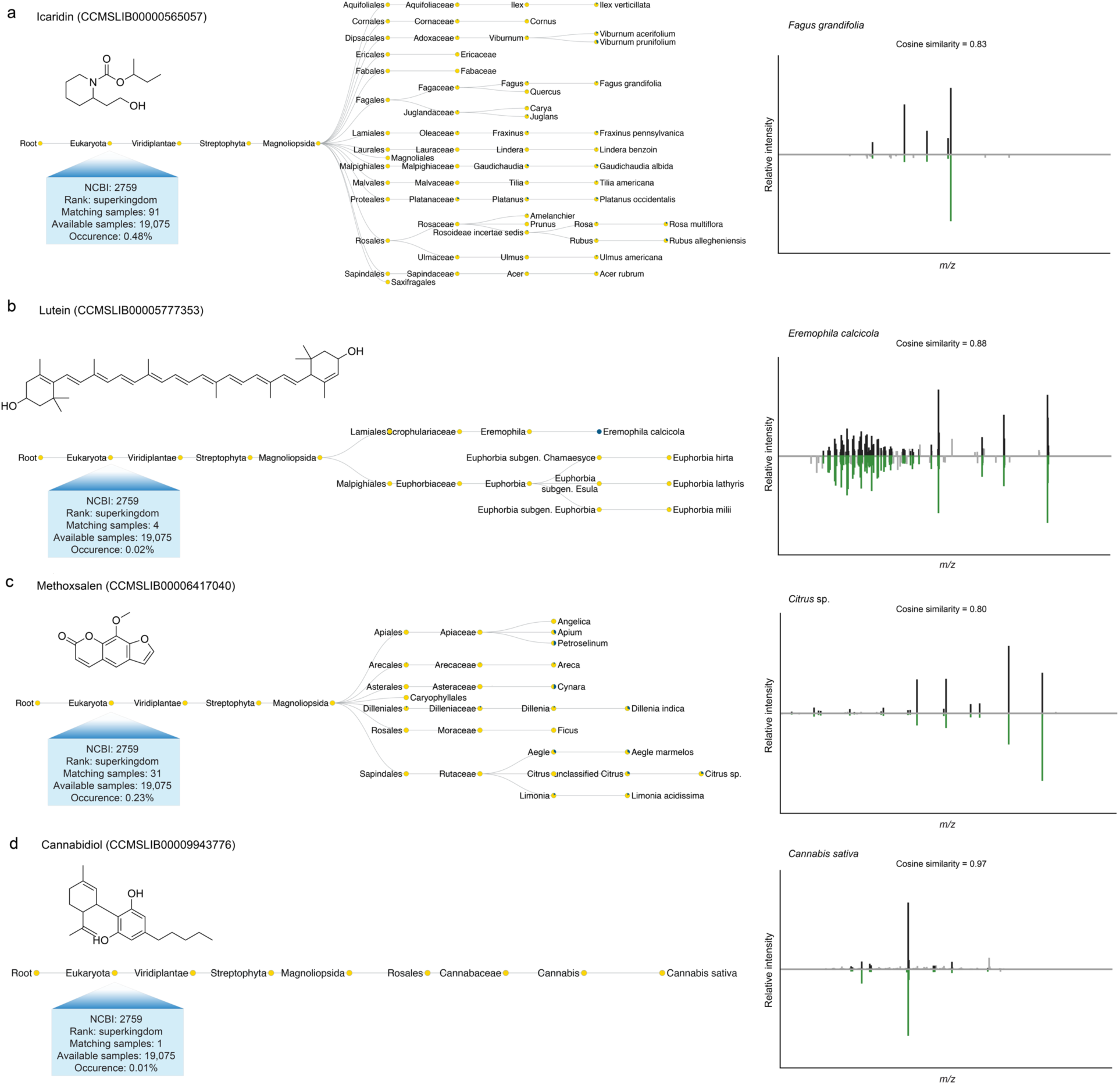
plantMASST examples of four rarely occurring plant-derived drugs. The plantMASST taxonomic trees for four drugs: a) icaridin, b) lutein, c) methoxsalen, and d) cannabidiol. Outputs showcase their occurrences in the reference datasets of plantMASST. The pie charts represent the percentage of matches within each taxonomic level detected against the plantMASST database. Blue indicates the percentage of samples with matches, while yellow without matches. The mirror plots show the matches between the representative MS/MS spectrum (top) of each drug and their closely matched MS/MS spectrum (bottom) in the plantMASST database. These are level 2 annotations according to the Metabolomics Standards Initiative^27^. a) icaridin, a piperidine-type alkaloid used as an insect repellent, was observed in Malpighiaceae plants known for their insecticidal and repellent properties. b) lutein, a carotenoid, was detected in various plant families, including Euphorbiaceae, which is a notable source of lutein. c) methoxsalen, a furanocoumarin known for causing photosensitization of the skin, was found in alternative plant sources with no prior references related to methoxsalen. d) cannabidiol, a cannabinoid used for various medical purposes, was observed in *Cannabis sativa*, known as the sole source of cannabidiol.

**Supplementary Figure S5.**
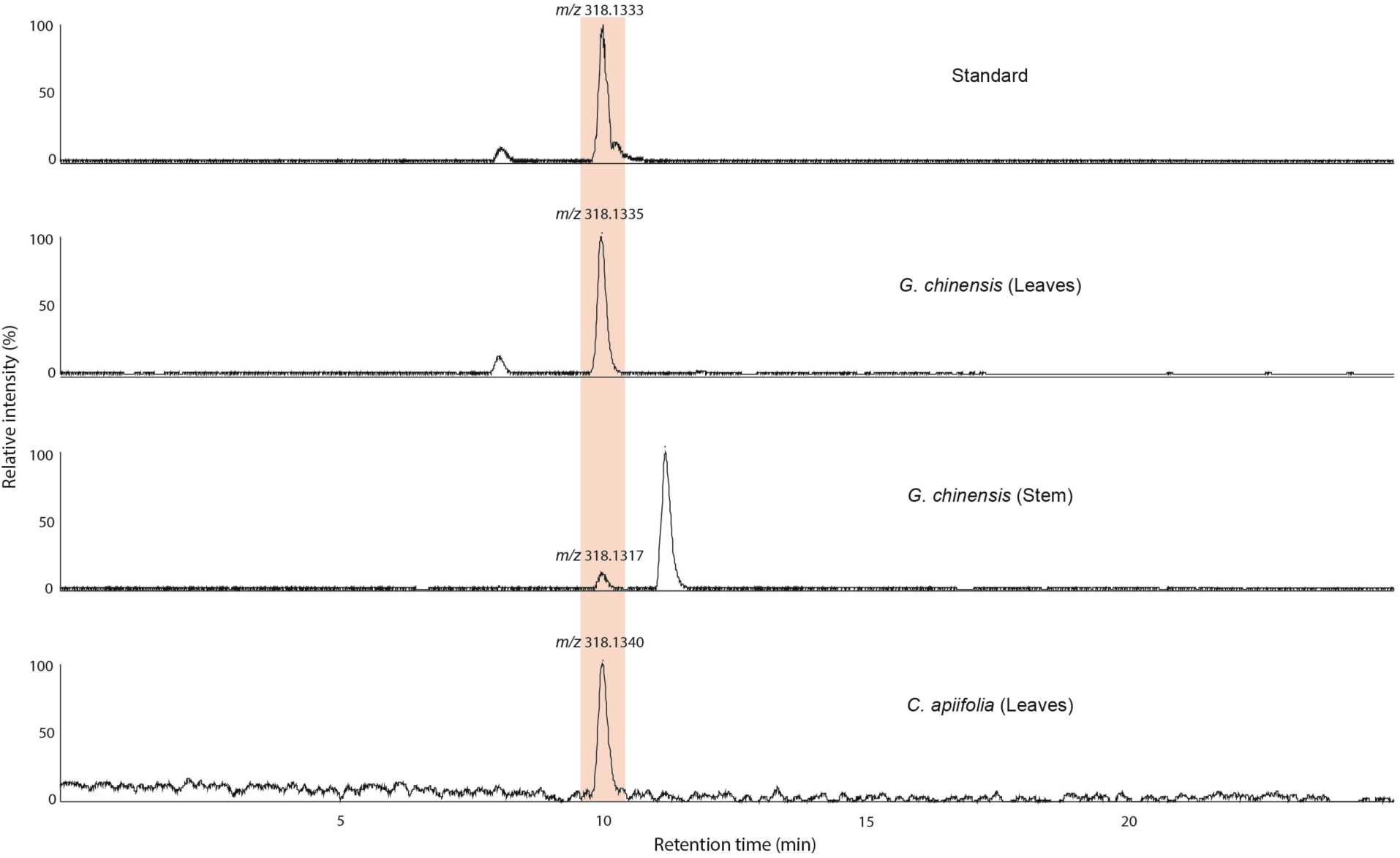
Retention time matching with piperlongumine. Retention time matching between piperlongumine (*m/z* 318, [M + H]^+^) commercial standards and *Gymnotheca chinensis* (leaves and stem) and *Clematis apiifolia* (leaves) extracts. Piperlongumine was not observed in the other extracts analyzed (*C. apiifolia* stems and roots). These are level 1 identifications according to the Metabolomics Standards Initiative^27^.

**Supplementary Figure S6.**
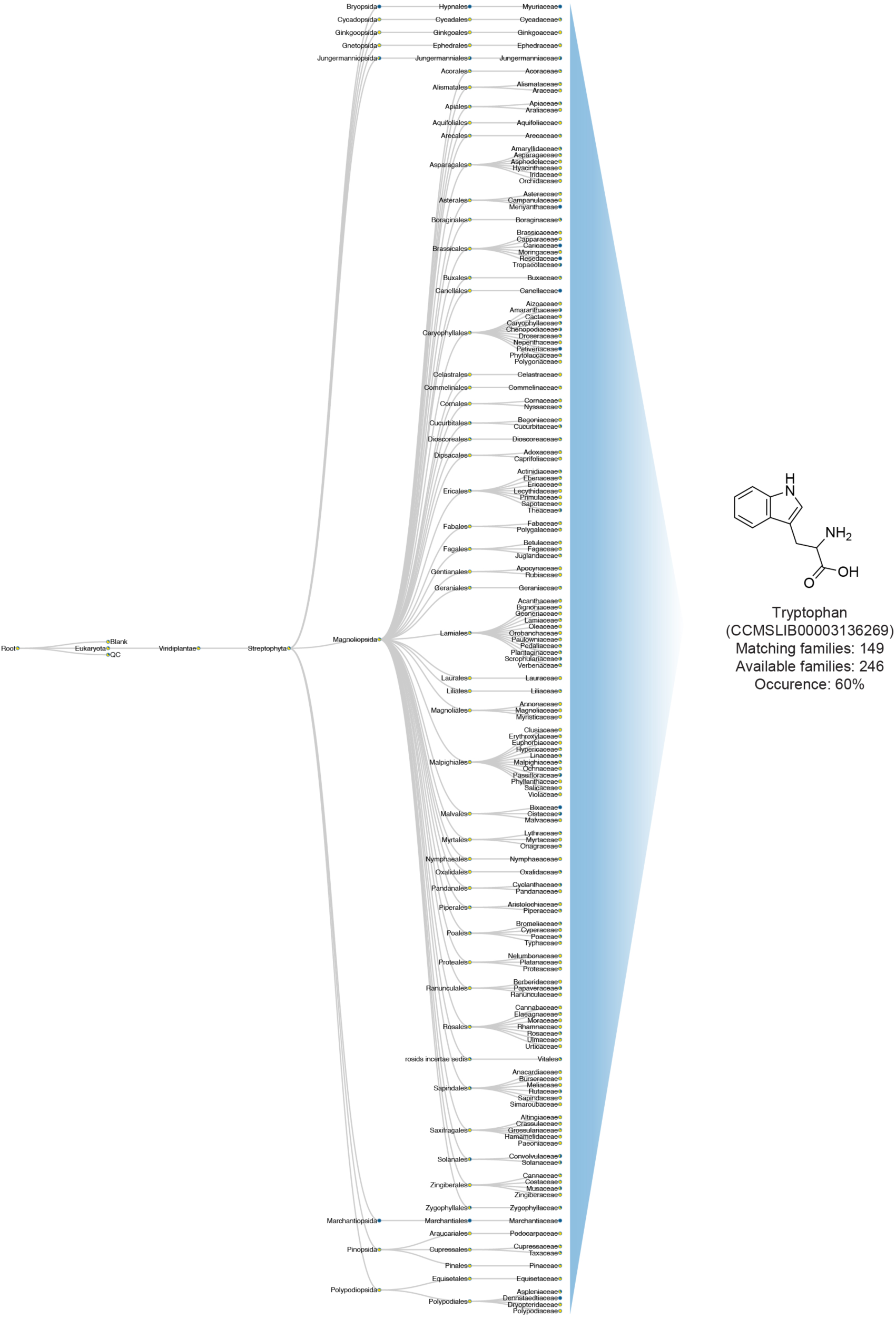
Taxonomic tree of tryptophan. Taxonomic tree of tryptophan (CCMSLIB00003136269) and its distribution abroad plant families according to the NCBI taxonomy.

